# Engineering butyrate-producing Lachnospiraceae to treat metabolic disease

**DOI:** 10.1101/2025.11.08.687059

**Authors:** Jack Arnold, Sandra McClure, Joshua Glazier, Recep E. Ahan, Sadiksha Shakya, David Villegas, Tracy Leidan Chen, Jay Fuerte-Stone, Rory McGann, Mark Mimee

## Abstract

Engineering native gut bacteria offers a route to persistent, programmable therapeutics, yet many dominant taxa remain genetically intractable. Lachnospiraceae are a prevalent and abundant family in the human gut microbiome, possessing metabolic functions generally associated with health^1^. Despite their promise as engineered live biotherapeutics, genetic manipulation of Lachnospiraceae remains challenging. Here, we develop a modular toolkit for Lachnospiraceae engineering, including constitutive and inducible expression and chromosomal integration systems. Applying this toolkit to the native commensal *Coprococcus comes*, we program secretion of the mammalian cytokine interleukin-22 (IL-22) in the mouse intestinal tract where it elicits ileal transcriptional responses consistent with cytokine signaling. In a mouse model of metabolic associated steatotic liver disease, IL-22–secreting *C. comes* improves glucose homeostasis and attenuates hepatic steatosis. This work demonstrates that a native Lachnospiraceae chassis can be genetically programmed to modulate host metabolic and immune physiology. The toolkit provides a generalizable foundation for Lachnospiraceae-derived microbiome therapeutics and for probing causal links between Lachnospiraceae gene programs and host phenotypes.

## Introduction

Microbiome-targeted therapeutics aim to edit the functional landscape of a microbial community to benefit the host. Supplementation of a deficient gut microbiome with beneficial functions natively encoded by microbial isolates or consortia has shown promise against infectious disease^2,3^, metabolic syndrome^4^, and psoriasis^5^ in human trials. With genetic engineering, therapeutic microbes can be further programmed with non-native metabolic functions^6,7^, heterologous protein secretion^8–11^, or controllable engraftment^7,12,13^ to program their behavior in the gut. Engineered live biotherapeutic products (eLBPs) have shown promise in treating metabolic diseases^6,7^, colitis^8,9^, and cancer^11,14,15^ in animal and early human trials, yet their clinical efficacy has often fallen short. While live biotherapeutics source microbial isolates native to the gut environment, most eLBPs rely on model microbes that represent a narrow subset of gut diversity and perform therapeutic activities only during gut transit. Employing a greater spectrum of gut microbes as chassis organisms could overcome limitations in colonization durability^16^ and provide access to a broader range of beneficial microbial metabolites.

New genetic methods have expanded access to previously intractable human-associated species^17–20^, including gut Clostridia. Members of Lachnospiraceae (previously *Clostridium* cluster XIVa), a family of obligate anaerobic Clostridia prominent in the human gut microbiome, are emerging as key mediators of host metabolism and health ^1,21–23^.

Lachnospiraceae play important roles in short chain fatty acid production^24^, conversion of secondary bile acid metabolites^22,25^, and colonization resistance^25,26^. In both inflammatory^27^ and metabolic diseases^28–30^, butyrate-producing Lachnospiraceae are often depleted in humans as compared to healthy controls. Furthermore, certain Lachnospiraceae species have been successfully employed as live biotherapeutics in mice, such as *Anaerostipes caccae* in food allergy^23,31^ and *Coprococcus eutactus* in metabolic associated steatotic liver disease (MASLD)^32^. Despite their prominence in the human microbiome, there are few genetic tools available for Lachnospiraceae engineering.

Here, we implemented a three-step roadmap to generate Lachnospiraceae-based eLBPs. First, we developed a foundational suite of molecular tools to enable genetic manipulation of multiple Lachnospiraceae species, including an *in vivo-*validated library of constitutive promoters that span a 10^6^ dynamic range, inducible expression systems, and a chromosomal integration strategy to increase the stability of transgene expression. Second, we defined a set of signal sequences to allow for *in situ* secretion of heterologous proteins. Finally, to establish viability as an eLBP, we programmed *Coprococcus comes*, a representative butyrate-producing Lachnospiraceae species, to secrete a therapeutic cytokine directly within the intestinal environment to protect against steatosis in a mouse model of MASLD. This work outlines foundational contributions to expand the therapeutic potential of Lachnospiraceae and the repertoire of commensal microbes as chassis organisms for eLBPs.

## Results

### Programming gene expression in Lachnospiraceae

The design of eLBPs necessitates a robust set of genetic parts to control heterologous gene expression in the complex gut environment. To prototype genetic constructs in Lachnospiraceae species, we extensively optimized inter-species matings between *Escherichia coli* and target Lachnospiraceae to transfer genetic cargo on shuttle vectors (pMTL plasmids)^33^, due to their previously reported ability to broadly transfer genetic material to Clostridia, including Lachnospiraceae species^17,34^ (Fig. 1a). We first developed a small library of promoters derived from common housekeeping genes (elongation factor EF-Tu and housekeeping sigma factor σ^70^) to drive expression of a NanoLuc reporter gene in taxonomically diverse Lachnospiraceae with a range of SCFA production capacity and aerotolerance (Extended Data Fig. 1a-b).

**Figure 1.**
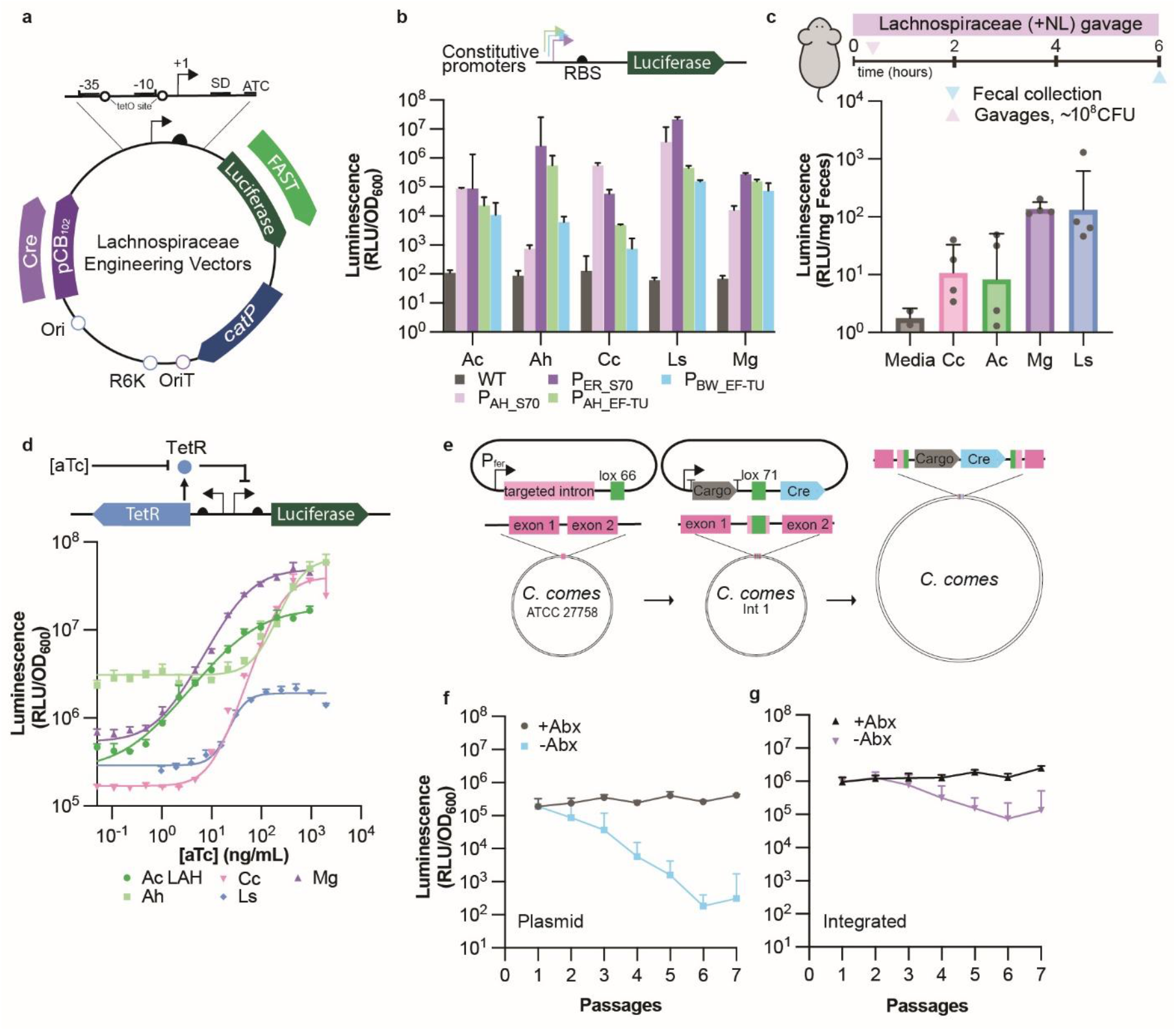
Genetic parts and integration toolkit for Lachnospiraceae engineering. (a) Schematic of Lachnospiraceae engineering toolkit derived from a vector-based platform. (b) Constitutive promoter screen across five Lachnospiraceae species; *A. caccae* (Ac), *A. hadrus* (Ah), *C. comes* (Cc), *L. symbiosum* (Ls), and *M. gnavus* (Mg). Promoters represent housekeeping genes σ70 from *A. hadrus* and *A. rectalis*, EF-Tu from *A. hadrus* and *B. wexlerae*. (c) Fecal bioluminescence 6 h after oral gavage with NanoLuc producing strains controlled by promoter P_AhS70_ (n=4). (d) Dose–response of TetR-regulated promoters to anhydrotetracycline (aTc), showing NanoLuc (NL) activity in Ac, Cc, Mg, Ah, and Ls. (e) Schematic of the two-step integration system for Lachnospiraceae genome engineering. A preliminary vector inserts a *lox66* landing pad using a retargeted type-II intron. Subsequent recombination with a cargo-bearing plasmid specifically inserts into the prepared site. (f, g) Maintenance of NL activity from plasmid or integrated constructs, with and without thiamphenicol selection. Data are presented as geometric mean ± geometric s.d.

NanoLuc reporters produced robust luminescence in type strains of *A. caccae, Anaerostipes hadrus, C. comes, Lachnoclostridium symbiosum*, and *Mediterranibacter gnavus* (Fig. 1b), as well as a human clinical isolate of *A. caccae LAHUC*^31^ (Extended Data Fig. 2a). The strong promoter P_AhS70_ also supported expression of a Fluorescence-Activating and Absorption-Shifting Tag (FAST), enabling interchangeable reporter proteins for flow cytometry and imaging applications (Extended Data 2b)^35^. Additionally, the strong P_AhS70_ promoter drove robust NanoLuc production compared to media controls in fecal pellets collected from specific pathogen free (SPF) mice after a single gavage (Fig. 1c), suggesting that engineered strains can deliver heterologous protein to the mammalian gut during transit.

Next, we adapted inducible systems in Lachnospiraceae species, allowing for exogenous control of genetic circuitry towards future therapeutic eLBP applications. Borrowing inducible systems developed for the study of *Clostidioides difficile* pathogenesis^36,37^, we first tested the efficacy of the xylose-inducible XylR repressor-P_xyl_ system (Extended Data Fig. 2d), the aTc-inducible TetR-P_tet_ system (Fig. 1d), and a synthetic TetR-P_tet_ system (Extended Data Fig. 2e) for dose-dependent induction of NanoLuc in multiple Lachnospiraceae species. Kinetics favored the TetR system, with NanoLuc peaking 3-6h post-induction, whereas xylose responses were slower (Extended Data 2f).

To improve the stability of genetic cargo and minimize the risk of horizontal gene transfer in the gut microbiome, we leveraged the *Lactococcus lactis* LtrA Group II intron^38,39^ modified with a *lox66* site to insert a landing pad into a fixed genomic locus with the *C. comes* chromosome (Fig. 1e and Extended Data Fig. 2g). Introduction of a non-replicative plasmid with a cognate *lox71* site and Cre recombinase into the landing pad strain led to engineered strains with cargo genes stably inserted in their genome^39,40^ (Fig. 1e). Chromosomal integrants demonstrated increased cargo stability during passaging in antibiotic-free media (Fig. 1f-g) with comparable expression levels. Overall, we describe a genetic toolkit for engineering heterologous gene expression in Lachnospiraceae species.

### Transcriptomics-informed expansion of the Lachnospiraceae expression toolkit

Next, we sought to expand the basic Lachnospiraceae engineering toolkit to include additional systems with a greater range of expression levels and conditional activity in culture broth or the mouse gut (Fig. 2a). To guide promoter library design, we performed differential expression analysis between Lachnospiraceae species (*C. comes, A. caccae*, and *M. gnavus*) monocultured *in vitro* or in the context of a 15-member defined microbial community in gnotobiotic mice (Fig. 2b, Extended Data 3a, and Supplementary Tables S6-8). Expectedly, Lachnospiraceae adapted to gut colonization through alterations in catabolic strategies, including carbon and amino acid utilization, fatty acid biosynthesis, and interbacterial competition (prophage, Type VII secretion) (Extended Data Fig. 3b-g). Using computational promoter-ID and differential expression analyses^41^, we identified 25 *C. comes* promoter-RBS candidates spanning a broad predicted dynamic range with condition biases (*in vivo*-biased, *in vitro*-biased, or both). The resulting *C. comes* library spanned ∼10^6^-fold range in *in vitro* gene expression (Fig. 2a-c and Extended Data 4b). Three promoters, representative of each class (P_3185_-*in vitro*, P_15920_-*in vivo*, P_4280_-both; Extended Data Fig. 4a) were functionally assessed in mice administered thiamphenicol to allow for durable gut colonization (Fig. 2e). Over the course of one week, P_4280_-NanoLuc engineered *C. comes* colonized mice exhibited greater luminescence in recovered fecal pellets, followed by *in vivo*-biased P_15920_, while *in* vitro-biased P_3185_ was barely detectable (Fig. 2d-e and Extended Data 4c). However, P_15915_ was strongly induced *in vivo*, as compared to either P_4280_ or P_3185_ (Fig. 2d-e). P_4280_ was further modularized through discovery of a minimal promoter motif and RBS modification (Extended Data Fig. 2c).

**Figure 2.**
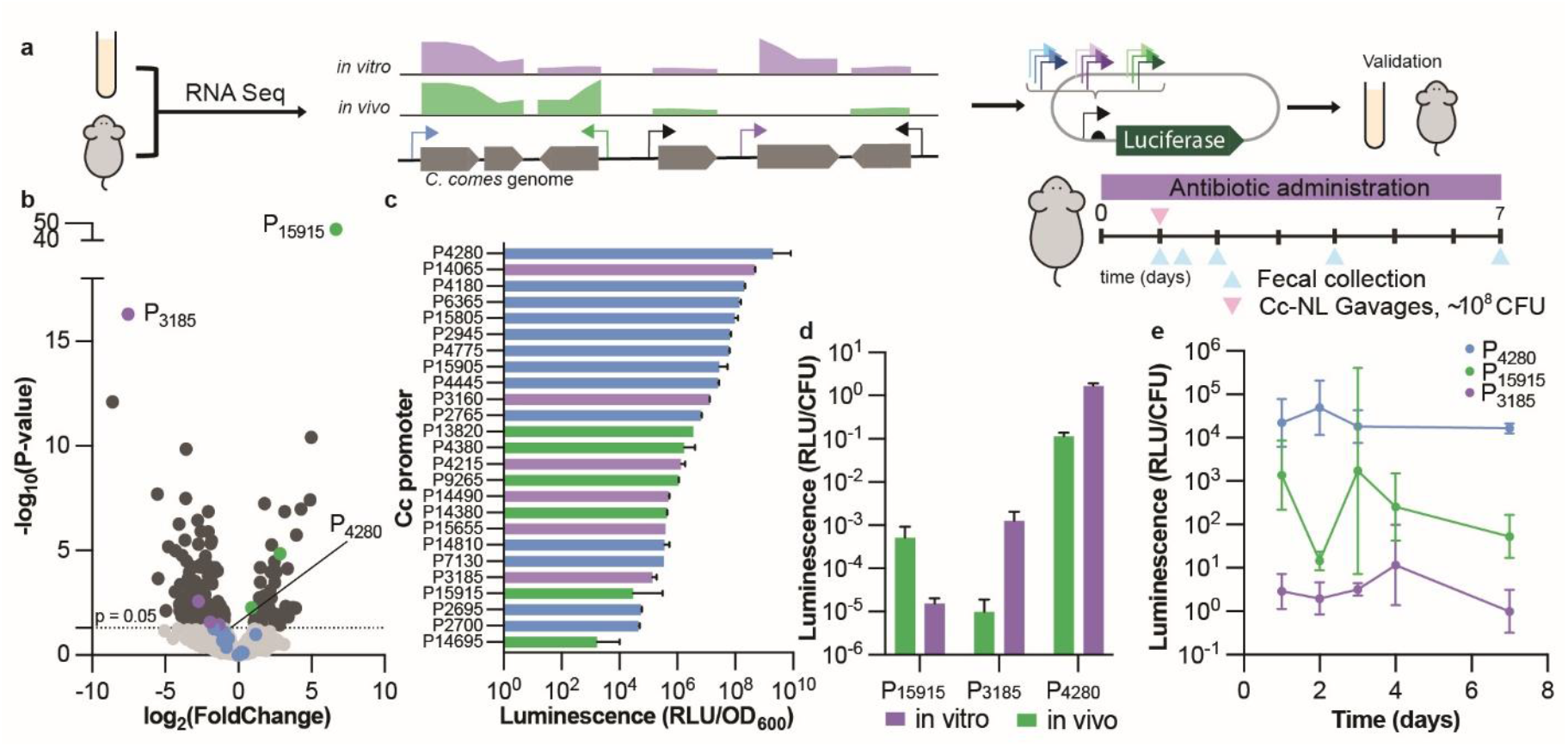
Transcriptomics-informed design of gut-active expression systems in Lachnospiraceae. (a) Schematic of the promoter-library discovery pipeline: starting from bacterial cultures or murine samples, RNA-seq was used to identify differentially expressed genes; upstream regions were extracted as candidate promoters, cloned upstream of a NanoLuc (NL) reporter, and tested *in vitro* and *in vivo*. (b) Volcano plot showing differential expression between *in vivo* and *in vitro* conditions for high expressing genes. Green: upregulated *in vivo*; purple: upregulated *in vitro*; blue: expressed in both conditions. (c) NanoLuc activity *in vitro* of RNA-seq-based promoter candidates in broth culture. (d) Comparison luciferase activity of selected promoter candidates P_15915_ (up-regulated *in vivo*), P_3185_ (up-regulated *in vitro*), and P_4280_ (highly expressed in both environmental conditions) measured either during mid-log growth (t = 3-5 hours, *in vitro*) or from Day 7 fecal pellets *(in vivo)*. (e) Longitudinal fecal bioluminescence from mice gavaged with *C. comes* expressing NL under promoters with predicted environment-specific activity monitored for 7 days. Drinking water contained thiamphenicol (15 μg ml−^1^) for plasmid maintenance. Points show geometric mean ± geometric s.d. *n* = 4 per group.

Together, these results establish a condition-aware native promoter set for *C. comes* and underscore the importance of environmental context in promoter performance.

### Developing secretion systems for Lachnospiraceae

To harness Lachnospiraceae as efficient eLBPs, we next sought to establish programmable secretion of heterologous proteins. Efficient secretion can maximize on-target delivery, while other strategies often introduce off-target effects and limit treatment longevity^42^. Guided by secretion systems from other Gram-positive bacteria^8^, we pursued a two-pronged discovery strategy combining proteomics and metagenomic mining (Fig. 3a). To discover Lachnospiraceae secreted proteins, we performed proteomics analysis of *M. gnavus* culture supernatant as compared to cellular lysate (Extended Data Fig. 5a, Supplementary Table S9-10). In parallel, we computationally predicted signal peptides in the *M. gnavus* genome using SignalP^43^ and generated a synthetic predicted signal peptide sequence from the consensus signal peptide motif (Extended Data Fig. 5b). These proteomically or computationally predicted signal sequences and signal sequences from known secreted proteins^44,45^ were cloned upstream of the NanoLuc reporter gene and the resultant library was introduced into *C. comes*. Most signal sequences increased extracellular NanoLuc relative to the no signal sequence control (Fig. 3a and Extended Data Fig. 5c). Interesting, the fully synthetic peptide REA14 exhibited >99% of total NanoLuc localized to the supernatant (Extended Data Fig. 5c), indicating highly preferential export. Additionally, we developed a surface display system based on the sortase-attached superantigen from *M. gnavus*, IbpA^45^ for cell adhesion or programmed localization applications^46^. We generated a series of truncated IbpA variants that maintain the N-terminal signal peptide, lack all superantigen domains, and include the C-terminal sortase motif. Using a trypsin sensitivity assay, we determined that the variant IbpA_379-607_ could efficiently display NanoLuc extracellularly in *C. comes* (Extended Data Fig. 5d). Together, our constructs establish a functional set of secretion signals for exporting or displaying protein cargoes in Lachnospiraceae.

**Figure 3.**
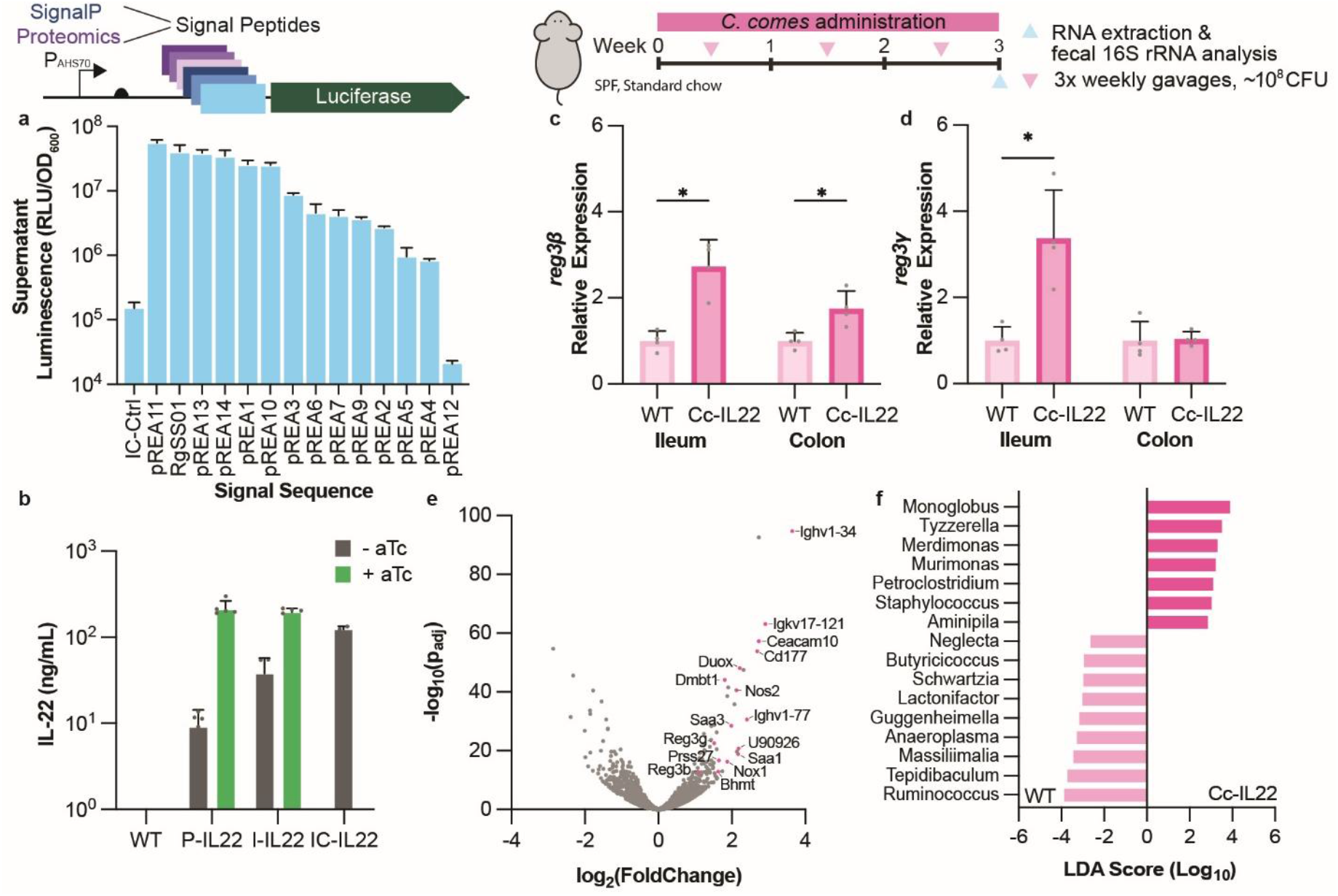
Engineered *Coprococcus comes* can modulate ileal gene expression in conventional mice. (a) Schematic of signal peptide discovery from proteomic analysis of supernatant vs cellular fractions and *in silico* mining from annotated ORFs. NL activity measured in *C. comes* supernatant fractions. (b) *In vitro* secreted murine IL-22 (mIL-22) production in *C. comes* (Cc) under an aTc inducible promoter, comparing expression from a plasmid (P), integrated construct (I), or integrated construct under a constitutive promoter, P_BW-EF-TU_ (IC). (c, d) Quantitative PCR analysis of host Reg3b and Reg3g expression in the ileum and colon after 3 weeks of bacterial administration in conventional, chow-fed mice. (e) Differential gene expression from host ileal tissue in mice administered WT Cc or Cc-IL22. (f) Genus-level taxonomic changes between WT Cc and Cc-IL22 administration at experimental endpoint. Error bars represent standard deviation. *, P < 0.05 (t-test). *n* = 4 mice per group.

### *C. comes*-produced mIL-22 alters host gene expression and microbiota composition

To demonstrate the therapeutic viability of engineered Lachnospiraceae, we modified *C. comes* to secrete IL-22 as a treatment for steatotic liver disease. Due to its consistent depletion in human studies of MASLD^28,29^ and preclinical work showing *Coprococcus*-mediated protection in mouse models of obesity^32^, we reasoned that *C. comes* could serve as a candidate chassis for an eLBP to treat metabolic disease. Additionally, we benchmarked engraftment and payload expression across candidate Lachnospiraceae species in mice. *C. comes* exhibited high intestinal reporter gene expression in chow and Western diet (WD)-fed mice and reliable dosing kinetics (Extended Data Fig. 5e-f). As a therapeutic payload, interleukin-22 (IL-22) is attractive for its capacity to support epithelial barrier integrity during gut and liver inflammation^42,47,48^ and to limit macromolecule uptake into circulation^49–51^. We introduced an RG_SS01_-murine IL-22 (mIL-22) fusion protein into *C. comes* (Cc-IL22) and mIL-22 was detected in culture supernatants under constitutive or aTc-inducible control and after genomic integration, demonstrating flexible expression formats (Fig. 3b). Peak titers were consistent with prior reports of recombinant mIL-22 with preclinical efficacy when delivered orally to mice^42,52^. Moreover, *C. comes*-secreted mIL-22 displayed comparable bioactivity to commercial recombinant mIL-22 in a STAT3-phosphorylation assay using human colonic epithelial HT-29-MTX cells (Extended Data Fig. 5g).

To demonstrate *in vivo* bioactivity, we treated SPF mice with either wild-type *C. comes* or Cc-IL22 (integrated mIL-22 under a constitutive promoter) by oral gavage three times weekly for three weeks. Cc-IL22 treatment led to *reg3b* and *reg3g* induction in ileal tissue, and to a lesser extent in the colon (Fig. 3c-d), consistent with the predominantly ileal activity of IL-22 on epithelial cells^48^. Bulk transcriptomic analysis of ileal tissue supported this signature: in Cc-IL-22–treated animals, we observed upregulation of *reg3* genes together with oxidative stress and antimicrobial programs, including *duox2, nox1, nos2*, and immune activation markers (*ighv1-34, igkv17-121, saa1*) (Fig. 3e and Extended Data Fig. 6a). Interestingly, Cc-IL22 elicited a transcriptional signature reminiscent of upregulated Th17 cell responses to segmented filamentous bacteria^53^, further indicating that *C. comes* produced mIL-22 affects phenotypic changes in a murine host.

Taxonomic profiling of the gut microbiome detected compositional changes due to Cc-IL22 treatment, including enrichment of *Monoglobus, Tyzzerella, Murimonas, Petroclostridium, Staphylococcus* in Cc-IL-22–treated mice, and *Ruminococcus, Tepidibaculum, Massiliimalia* in wild-type *C. comes*-treated mice (Fig. 3f). We also noted time-dependent decreases in Akkermansiaceae and Ruminococcaceae over the duration of *C. comes* treatment as compared to baseline (Extended Data Fig. 6b). Together, these data indicate that IL-22 delivery from a native Lachnospiraceae chassis is bioactive *in vivo*, engages epithelial defense pathways, and is accompanied by measurable shifts in the intestinal community beyond the intrinsic metabolites of *C. comes*.

### IL-22–secreting *C. comes* attenuates steatosis and hepatocellular injury in a diet-induced model of MASLD

Building on the observation that IL-22 secretion activates epithelial defense and restoration pathways^48^ and on prior reports of protective IL-22 activity in metabolic disease^49,50^, we tested Cc-IL22 in a mouse model of MASLD. Modifying an established diet-induced metabolic steatosis model^49,54^, we provided mice a WD and 30% fructose-supplemented water (to exacerbate liver fat accumulation) for 8 weeks to establish disease prior to *C. comes* treatment (Fig. 4a). During the treatment period, mice were split into separate cohorts and administered bacterial growth media, wild-type *C. comes*, or Cc-IL22. Bodyweight trajectories diverged after treatment onset; both *C. comes* groups gained weight more slowly than WD controls, with the largest attenuation in the Cc-IL22 cohort (Fig. 4b-c). As an intermediate metabolic check, fasting glucose showed a modest, non-significant reduction in treatment groups (Extended Data Fig. 7a). After 7 weeks of bacterial administration, glucose tolerance tests (GTTs) revealed improved glucose clearance in mice given Cc-IL22 as compared to vehicle and wild-type *C. comes*-treated mice (Fig. 4d-e). The subsequent insulin tolerance test (ITT) showed significantly greater insulin sensitivity in the Cc-IL22 cohort than in wild-type or vehicle groups (Fig.4f-g). These results, together with compositional analyses (Extended Fig. 7b-h), establish payload-dependent efficacy from cytokine delivery, as opposed to relative changes in taxonomic or metabolomic profile of the microbiome.

**Figure 4.**
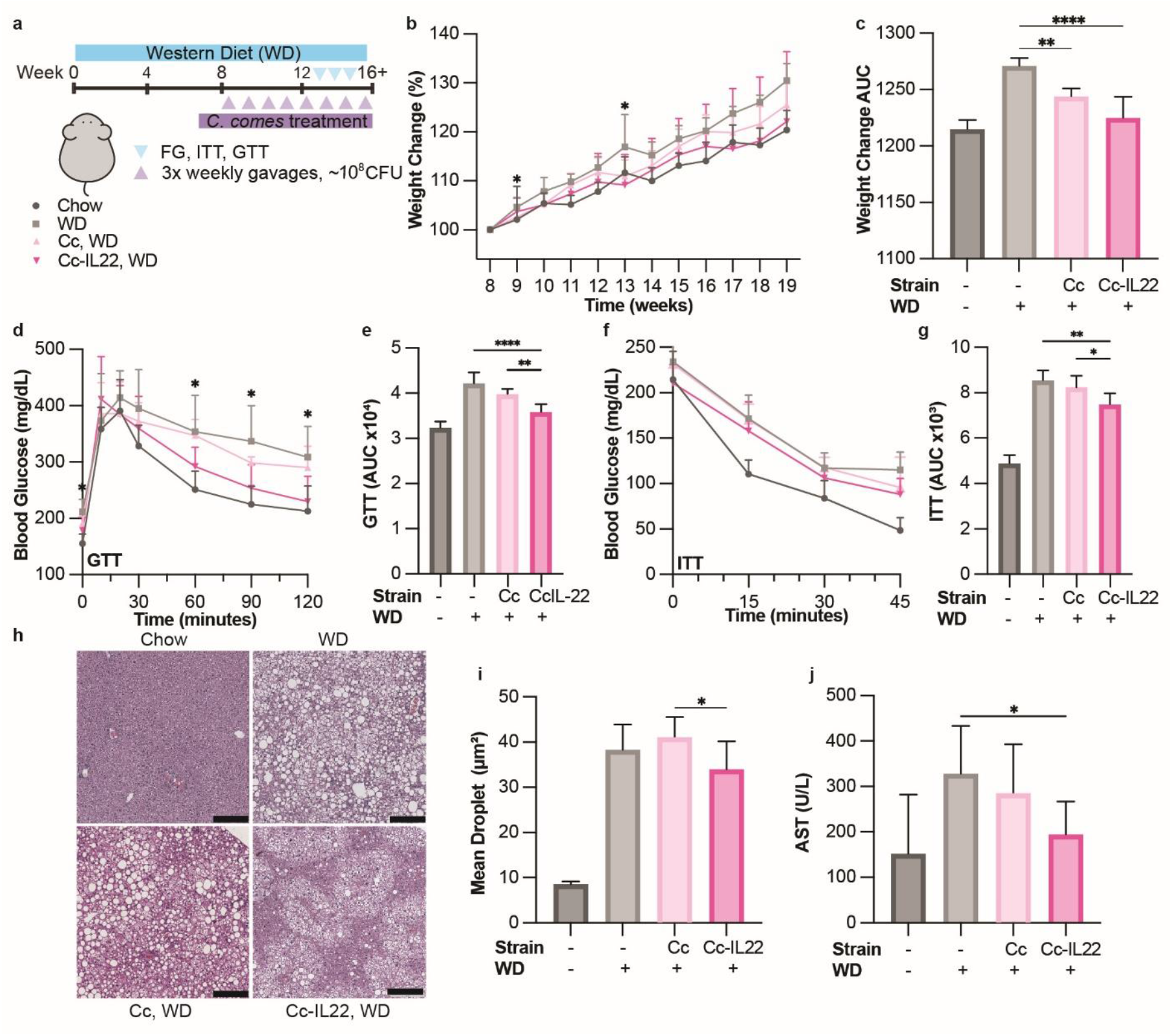
Engineered *Coprococcus comes* improves metabolic outcomes in a mouse model of metabolic associated steatotic liver disease. (a) Schematic of the Western diet (WD, 42% calories from fat, 30% fructose water)–induced steatosis model. *n = 10* per group for Chow, *C. comes* (Cc), and *C. comes–IL22* (Cc-IL22); *n = 5* for the untreated WD group. Mice in the Cc and Cc-IL22 groups received ∼10^8^ CFU per mouse three times weekly. (b) Body weight monitored throughout treatment; weight gain is shown as change from baseline. *P* < 0.05 between Cc-IL22 and WD groups. (c) Time integrated area under the curve (AUC) for all Chow- and WD-fed groups. (d, e) Glucose tolerance test (GTT) and corresponding AUC. *, P < 0.05 between Cc-IL22 and HFD groups. (f, g) Insulin tolerance test and AUC. (h) Representative H&E-stained liver sections, illustrating mean fat droplet size for each group. (i) Quantification of mean fat droplet size across all liver lobes. (j) Serum aspartate aminotransferase (AST) levels. (k) Error bars represent mean ± s.d. Statistical significance determined by one-way ANOVA:*, P < 0.05; **, P < 0.01; ***, P<0.001; ****, P<0.0001.

MASLD is characterized by hepatocellular lipid accumulation and injury, processes that IL-22 can mitigate by strengthening epithelial barrier function and limiting macromolecule flux to the liver. In our WD/30% fructose model, mice receiving Cc-IL22 showed slightly lowered liver mass relative to controls (Extended Fig. 8a). However, histological analysis revealed attenuated steatosis as mean lipid-droplet area was reduced and the size distribution shifted away from large droplets, with a selective decrease in the largest size bins while small–medium droplets remained largely unchanged (Fig. 4h-i, Extended Data Fig. 8b-d). Biochemical indices were concordant with these morphological improvements; aspartate aminotransferase (AST) fell significantly, with alanine aminotransferase (ALT) and hepatic triglycerides modestly reduced but not significantly (Fig. 4j, Extended Data Fig. 8e-f). Taken together, the significant AST reduction and decreased large-droplet burden indicate that IL-22 delivery from a native Lachnospiraceae chassis mitigates hepatic injury and steatosis in diet-induced MASLD.

## Discussion

As stable gut colonizers that produce beneficial metabolites, commensal microbes are optimal chassis organisms for eLBPs. Here, we described a robust genetic toolkit for the manipulation of Lachnospiraceae species, a prevalent and abundant family of butyrate-producing commensals. We identified promoter systems that enable *in vitro* and *in vivo* control of gene expression in an array of Lachnospiraceae species. Native promoter–RBS parts derived from *C. comes* transcriptomes span a broad dynamic range and exhibit condition bias, enabling context-aware control. The resulting toolbox also established inducible control of gene expression in Lachnospiraceae, enabling fine-tuned exogenous control of gene expression and establishing a foundation for responding to environmental stimuli ^18,20^. By co-opting an intron-based landing pad system ^39,40^, we expanded previous type-II intron-based Lachnospiraceae engineering from loss-of-function^17^ to gain-of-function applications, improving the stability of genetic circuitry in Lachnospiraceae.

Beyond expression control, we describe methods to expand the capacity of Lachnospiraceae to produce biologically relevant protein targets. Using proteomic and computational approaches, we developed a library of signal sequences to enable Lachnospiraceae to facilitate intestinal delivery of recombinant protein targets. As a proof-of-concept, we engineered *C. comes* to deliver mIL-22 to the mouse ileum, where it induced upregulation of gut immunity and altered the gut microbiome. When tested in a diet-induced mouse model of MASLD, we demonstrated Cc-IL22 can mitigate weight gain, insulin resistance, and hepatic steatosis. Future challenges in translating Lachnospiraceae-based therapies will address their fastidious growth requirements and sensitivity to oxygen. Directed evolution and syntrophic interactions can be used to improve the therapeutic viability of similar butyrate-producing taxa^55^. Altogether, these results present native Lachnospiraceae as a tractable taxa that can be co-opted for live biotherapeutic purposes.

## Methods

### Strains and Culture Conditions

All strains used are listed in Supplementary Table S1 along with their antibiotic sensitivity in Supplementary Table S5. Unless otherwise noted, *E. coli* strains were grown aerobically at 37C in LB broth while shaking at 250 RPM (BD Difco) or on plates (agar 15 g/L) (Fisher Scientific). Strains of Lachnospiraceae bacteria were grown anaerobically (anaerobic chamber from COY Systems) at 37C in BHIS media: 37g Bacto BHI Premix, 5g Yeast extract (Gibco), 4ml Resazurin (Arcos Organics) at 250 μg/mL, 1 mL Tween80 (Sigma) before autoclaving, and 10 mL L-cysteine-HCl (EMD Millipore) at 50 μg/mL, 1 mL menadione (MP Biomedical) at 1 mg/mL, 10 mL Hemin (Sigma Aldrich) at 1 mg/mL, and 20 mL Trace Mineral Supplements (ATCC) were added after autoclaving.

Antibiotics and other supplements were provided at the following concentrations as noted: chloramphenicol – 12.5 μg/mL, kanamycin – 30 μg/mL (aerobic cultures) or 60 μg/mL (anaerobic cultures) (GoldBio), thiamphenicol – 5 μg/mL (RPI), anhydrous tetracycline – 100ng/mL (Themo Fisher Scientific). Dilutions of bacteria for CFU assays were performed in 1x phosphate buffered saline (Thermo Fisher Scientific) with or without 0.05% L-Cysteine-HCl (aerobic or anaerobic respectively).

### Plasmid Construction & Transformations

#### Constitutive promoter plasmids

Plasmids used in this work are listed in Supplementary Table S2 and genetic parts in Supplementary Table S4. To construct plasmid pJHA105, a 3-piece Gibson assembly was performed following standard procedures ^56^. The 167 bp region upstream of σ70 gene P_AHS70_ (locus: NAGOKMPI_02529) from *Anaerostipes hadrus* DSM 3319 was extracted via PCR as a putative promoter region and cloned upstream of a gBlock (IDT) of the NanoLuc coding sequence ^57^, which was first codon optimized for *Clostridium acetobutylicum* ATCC 824 using the IDT Codon Optimization Tool. Both parts were subsequently cloned into pMTL83151 vector _33_ into the *lacZα* fragment between BamHI and HindIII restriction endonuclease sites.

Plasmids containing other promoter regions were constructed *via* Gibson assembly by extracting promoter regions from *Blautia wexlerae* (P_BW-EFTU_), Anaerostipes hadrus (P_AH-EFTU_), *Eubacterium rectale* (P_ER-S70_), pRPF185 ^37,58^, (P_TetA_), or pIA33 ^36^ (P_Xyl_), and subsequently cloning these putative promoter regions upstream of optimized NanoLuc in pJHA105.

#### Inducible promoter plasmids

To construct plasmid pJHA272, a 2-piece Gibson assembly was performed following standard procedures ^56^. The TetR-P_TetA_ regulator-promoter region was extracted via PCR from plasmid pRPF185 ^58^ and cloned into pJHA105 (this work) replacing the P_AHS70_ promoter region.

To construct plasmids containing signal peptides (pJHA521-pREA14; this work), signal sequences were synthesized into primers (IDT) and cloned using site directed mutagenesis cloning techniques to insert <∼60 bp fragments upstream of the NanoLuc coding region. Full-length signal peptide fusions were cloned by extracting signal sequence-secreted peptide DNA from purified *Medditeranibacter gnavus* ATCC 29149 genomic DNA and performing Gibson assembly using standard techniques ^56^.

#### Plasmids containing type-II intron machinery

To construct plasmids containing type II intron machinery, a control vector (pJHA226) was created by 4-piece Gibson assembly. PCR was used to extract P_fer_ promoter and the *ltrA* intron-encoded reverse transcriptase from the plasmid pQint ^59^. The P_fer_ promoter was encoded upstream of a retargeted intron sequence (ClosTron ^60^) gBlock (IDT), followed by the *ltrA* gene and into the pJHA105 vector backbone replacing both P_AHS70_ and the NanoLuc coding sequence. To create a landing pad site, pJHA226 was digested with the restriction enzyme MulI (NEB) and a *lox66 attB* site was inserted to match *lox* positioning in the plasmid pQlox66 ^40^ using isothermal assembly ^56^ techniques construct plasmid pJHA275.

To target chromosomal regions of Lachnospiraceae species, retargeted intron sequences were ordered as gBlocks from IDT and cloned into pJHA275 (this work). Retargeted intron sequences were designed using the ClosTron design tool ^60^ using the Perutka method. This method was used to create plasmids for landing pad strain engineering including *C. comes*-targeting introns (pJHA787; this work).

To create *cre* recombinase-based insertion vectors, plasmid pJHA789 was constructed *via* 4-piece Gibson assembly using the origin RP4/RSK from pAT890 ^57^, the P_AHS70_ promoter, a gBlock containing codon optimized *cre* recombinase ^40^ (*Clostridium acetobutylicum* ATCC 824; IDT codon optimization tool) and the *lox71 attP* site with random spacers which was inserted downstream of the P_AHS70_ promoter, and the *catP* antibiotic resistance gene (pJHA105; (this work)).

An inducible NanoLuc was subsequently added to pJHA789 to create pJHA830 by 2-piece Gibson assembly of TetR-P_TetA_-NanoLuc flanked by terminator regions from pJHA272 into the spacer region downstream of the *lox71 attP* site.

Bioactive protein cargo (with or without signal sequences) was added to pJHA830 *via* Gibson assembly to replace the NanoLuc coding sequence.

### Bacterial Conjugation

To transfer genetic material from donor *E. coli* S17 λpir^−^ to Lachnospiraceae species, conjugative mating strategies were adapted from Sheridan *et al*. 2019 ^34^. *E. coli S17* λpir^−^ was chosen as a donor due to its suitability for most molecular cloning applications while maintaining conjugation machinery^57^. To determine suitable parameters for selection we first performed a mean inhibitory concentration (MIC) assay (Table S1) for selection in a Lachnospiraceae recipient and counterselection against the *E. coli* λpir^−^ donor. Prior to mating, plates containing supplemented Brain Heart Infusion agar (BHIS; BD) containing 0.05% cysteine-HCl as a reducing agent were added to the anaerobic chamber and allowed to reduce at least 4 hours for optimal mating conditions. Following successful construction of shuttle vectors, saturated overnight cultures in selective media of *E. coli* containing cargo DNA are diluted 1:100 in fresh selective media and grown for approximately 4 hours to reach OD_600_ of 0.4-0.6.

Concurrently, saturated overnight cultures of recipient Lachnospiraceae species are diluted 1:100 and grown for approximately 4 hours to reach OD_600_ of 0.4-0.6. Once reached, 1mL of donor *E. coli* culture is centrifuged for 5 minutes at 5000xg. The supernatant is discarded, and the donor pellet is transferred to an anaerobic chamber (COY Systems) where it is washed twice with anaerobic PBS containing 0.05% cysteine-HCl (centrifugation at 5000xg for 5 minutes). After washing, 250 μL of Lachnospiraceae recipient culture is used to resuspend the *E. coli* donor pellet and the entire suspension is carefully added to reduced BHIS plates in a single spot and rested at room temperature for at least 30 minutes to allow for the spot to dry.

Mating plates are then incubated anaerobically at 37C anaerobically for 18-24 hours. Following incubation, mating spots are scraped using sterile 1 μL inoculation loops into 500 μL of anaerobic PBS and suspended by aspiration. After a homogeneous suspension is achieved, 250 μL of the suspended mating is plated on reduced selective BHIS plates containing 5 μg/mL Thiamphenicol and 60 μg/mL Kanamycin and plates are incubated at 37C for 24-72 hours to allow colonies to appear. After colonies appear on selective plates, individual colonies are selected using a sterile inoculation loop and streaked onto fresh reduced selective BHIS plates to isolate single colonies. For conjugation using *Coprococcus comes ATCC 27758*, the “sticky” nature of the bacteria facilitated morphological differentiation from *E. coli* donor cells.

Isolation plates are incubated anaerobically at 37C for 24-72 hours and isolated colonies are grown in reduced selective BHIS media anaerobically at 37 C for 24-72 hours. Colony PCR with GOTaq2 (NEB) is used to confirm the successful transfer of genetic material.

### Luciferase Activity Measurements

To measure gene expression, NanoLuc was used as a reporter gene in Lachnospiraceae species. Cells were routinely cultured in media supplemented with antibiotics when necessary for until stationary phase was achieved (16-24 hours). Cells were then diluted 1:100 in fresh media (containing appropriate antibiotics or supplements as noted) unless otherwise indicated for 8-20 hours. For full-cell and supernatant expression measurements, equal volumes of cell culture and Nano-Glo working reagent (1:50; Nano-Glo Substrate: Nano-Glo buffer) (Promega) were added to a flat white 96 well plate (Costar) and luminescence was recorded (Tecan infinite 200).

### FAST Activity Measurements

As an alternative to NanoLuc, Fluorescence-Activating and Absorption-Shifting Tag (FAST) was used to quantify gene expression^35^. Cells were routinely cultured in media supplemented with antibiotics when necessary until stationary phase was achieved (16-24 hours). Cells were then diluted 1:100 in fresh media (containing appropriate antibiotics or supplements as noted) for 8-20 hours (unless otherwise indicated). Once logarithmic phase was reached, 1 mL of cells were harvested from liquid cultures (centrifugation 5000xg for 5 minutes). Cells were then resuspended and normalized to OD_600_ of 1 in either 1x PBS or 1x PBS containing 20 μM 4-hydroxy-3-methylbenzylidene-rhodanine (HBMR; Sigma). Fluorescence (excitation 485 nm; emission 535 nm) was measured via microplate reader using a flat bottom clear black microplate (Costar) containing 200 μL of normalized sample.

### Promoter library construction

#### Bacterial RNA-Seq Analysis

All steps involving RNA handling were performed with sterile filter pipette tips (Fisher Scientific) and RNase Away spray was used to clean surfaces. For *in vitro* sampling, Lachnospiraceae were grown in BHIS media overnight then subcultured 1:50 until mid-log growth, as measured by OD_600_. RNAProtect (Qiagen) was added to the cells per manufacturers protocols before RNAisolation. Total RNA was isolated from each cell type (in biological duplicate) and purified using RNAeasy RNA extraction kit (Qiagen). For *in vivo* sampling, a 15-member defined community that included *C. comes, M. gnavus*, and *A. caccae* was sampled 2 weeks post gavage in germ-free mice. At 2 weeks post gavage, mice (n = 3) were sacrificed to harvest cecal and colon contents. RNA from *in vivo* contents was isolated via acidic phenol:chloroform extraction (Thermo Fisher AM9720) and purified via Qiagen RNeasy columns and protocols^61^.

All RNA was then DNAse treated (TURBO) and submitted to the UChicago DFI Metagenomics Core for Illumina RNAseq Library Prep. Illumina compatible libraries were generated using NEBNext® Ultra™ II Directional RNA Library Prep Kit. Ribosomal RNA depletion (NEB) was used prior to library preparation. Libraries were sequenced using 2×150bp reads on NextSeq 1000. After sequencing reads were filtered for quality (FastQC), they were aligned to the respective reference genomes^62^ (Bowtie). Aligned reads counts were binned to reference annotations using featureCounts^63^ (Rsubread) and tested for differential expression (DESeq2)^64^.

Differential expression analysis directly compared the *in vitro* vs *in vivo* (cecal and colon) conditions, generating lists of significantly up- and down-regulated genes (adjusted p-value < 0.05 and log2 (fold change) either > 2 or < −2). Lachnospiraceae genomes were further annotated via PATRIC (BV_BRC) to add SEED Subsystems. Enrichment analysis was performed using the GSEA() function from the clusterProfiler^65^ package to search for subsystems significantly enriched (P < 0.05). All analysis was done in Rstudio and plots generated using the tidyverse R package.

#### RNA-Seq-based promoter library

For RNA-Seq-based promoter design, RNA was extracted, isolated, and analyzed as described below. Promoters were identified using promoter-id-from-rnaseq^41^ to find promoters of highly expressed genes *in vitro* and *in vivo*, and from differential expression analysis to find promoters specific to each condition. Promoters of various lengths (75 – 300 bp) were isolated by genomic DNA PCR and cloned into a NanoLuciferase expression plasmid (pJHA105) using Gibson and Golden Gate Assembly methods. Promoter sequences were confirmed by Sanger sequencing and then mated into *C. comes* according to previously mentioned methods.

*In vitro* promoter expression was determined by picking *C. comes* colonies into overnight BHIS cultures, growing overnight, and subculturing 1:100 to mid-log growth (3-5 hours) in pre-reduced BHIS media. Cultures were resuspended thoroughly before OD_600_ and RLU measurements.

Specific promoters were also plated on antibiotic BHIS plates to count Colony Forming Units (CFUs) at mid-log growth time points for RLU normalization.

### Induction curves

Inducible gene expression was measured by a NanoLuc luminescence assay. Briefly, Lachnospiraceae cells containing pJHA272 (this work) were grown at 37C in BHIS media containing the appropriate antibiotics until reaching stationary phase (16-24h). Cells were diluted 1:100 into fresh media containing aTc (10-fold dilutions ranging from 2 μg/mL aTc to 1.95 ng/mL with an aTc-free control) in a 96-well plate and grown for 24 hours at 37C. Following luminescence assay measurements, response curves were fit to a Hill function: Y = (R_max_X^n^)/(K^n^ + X^n^) + B, where X is the input, Y is the output, R_max_ is the maximum luminescence above the baseline, n is the Hill coefficient, K is the threshold (EC_50_), and B is baseline luminescence.

### Type-II intron design

Type-II introns were used to insert approximately 1Kb of DNA into Lachnospiraceae genomes, containing integrative *lox att* sites when denoted. ClosTron design tools ^60^ were used to determine optimal exon targeting sequences, preferentially choosing scores >5.0 when possible. Plasmids containing retargeted introns, *ltrA*, and *lox* landing pad sites were assembled as described above.

### Integration conjugation and verification

Conjugation of intron vectors followed standard conjugation procedures (above). Following verification, remaining plasmid (containing antibiotic resistance marker genes) was cured by serial passaging in reduced antibiotic-free BHIS broth and individual colonies were streaked on both reduced selective BHIS plates containing 5 μg/mL Thiamphenicol and 60 μg/mL Kanamycin and reduced antibiotic-free BHIS plates to confirm plasmid loss.

Integration of type II introns was initially verified *via* colony PCR using primers (JHA.L12 and JHA.L13, Supplementary Table S3) flanking the insertion region using GOTaq2 polymerase to identify the ∼1Kb increase in band size. Following identification of potential insertion events, Sanger sequencing was used to confirm integration fidelity and orientation.

Recombination events between *lox* landing pad sites and *cre/lox* cargo vectors were screened for with primers flanking the resulting *lox72* site following *cre/lox* recombination events (primers facing *lox71* site with homology to the *cre* vector and facing the *lox66* site with homology to the type II intron vector respectively (JHA.L13 and JHA.L36, Supplementary Table S3). Sanger sequencing was used to confirm integration fidelity.

### Plasmid and integrated cargo maintenance assays

All Lachnospiraceae culturing steps were performed in an anaerobic chamber. NanoLuc encoded on either a replicative plasmid or integrative vector was transferred to Lachnospiraceae following standard conjugation procedures (above). Cells were grown to saturation in triplicate before measuring NanoLuc gene expression. Cultures were then diluted 1:100 in fresh media with or without selective antibiotics and grown to saturation before measuring NanoLuc gene expression. This process is then repeated for the duration of the maintenance assay.

### Secretome preparation and analysis

For secretome analysis, *M. gnavus* cells were grown at 37C in heme-free YPG media (Anaerobic Systems) until saturation. Supernatant was collected via centrifugation (5000xg, 20 minutes) and concentrated 100-fold using concentrator filters (Amicon). Cell pellets were washed with 1X PBS and lysed (SDS lysis buffer (5% SDS, 50mM TEAB, pH 7.55). Samples were prepared for MS analysis by the University of Chicago proteomics core by reduction, alkylation, and a trypsin digestion followed by a C18 stage tip clean up. MS samples were run in duplicate and detected proteins were identified using custom ORFs^66^.

A volcano plot was drawn to compare proteins that appeared both in the supernatant and cell lysate. Proteins significantly enriched in the supernatant fraction compared to cell lysate (< −2 indicates a 2-fold change enrichment, p < 0.05) were selected and their signal peptides were tested in downstream NanoLuc assays to assess secretion potential (Extended Data Fig. 5a).

### Signal peptide library prediction and construction

#### SignalP cleavage analysis and reporter sequence construction

Annotated CDS from *M. gnavus* was translated and signal peptides were predicted using the SignalP 6.0^43^ webserver. Predicted signal peptides were selected for testing based on annotated functionality, and peptides corresponding to proteins secreted *via* the Sec pathway were preferentially selected. Three signal sequences predicted by SignalP 6.0 (SS) and their corresponding proteins (SP) were selected from *M. gnavus* for analysis.

Each SS reporter construct consists of an ATG start codon, the predicted SS sequence from SignalP inclusive of predicted the cleavage site immediately upstream of a NanoLuc sequence lacking the start codon. SP reporter constructs contain full protein sequences corresponding to predicted signal peptides and a Gly-Ser linker (GGSGGGSGGGSG) immediately upstream of a NanoLuc sequence containing the start codon. Reporter constructs were cloned into an aTc-inducible vector (pJHA272; this work) replacing the existing NanoLuc sequence.

#### Synthetic signal peptide prediction

A single motif of all *M. gnavus* predicted signal peptides from SignalP 6.0 was generated using Glam2 from the MEME-suite web server^67,68^. Based on the predicted motif, a single amino acid change was made (G10L) to maintain a positively charged N-terminal region and the resulting SS was cloned into the reporter vector as described above.

#### Signaling of known secreted proteins

Using SignalP 6.0 ^43^, signal peptide sequences were determined for previously described intramolecular *trans*-sialidase (*nanH*; RUMGNA_02694)^44^ or immunoglobulin binding protein (*ipbA*; WP_105084811.1)^45^. Signal sequences for these proteins were separately cloned into expression vectors as described above. In addition, a signal peptide construct for *nanH* was extracted *via* PCR from the *M. gnavus* genome for characterization with the secreted NanoLuc assay. Since IbpA has a sortase-like anchoring motif, it was not used as a whole protein fusion for secretion experiments^45^.

#### Surface anchored protein cargos

NanoLuc was fused downstream of the predicted signal sequence for IbpA^45^ and truncations of IbpA containing a sortase binding motif (LPXTG) were added downstream of the NanoLuc CDS using a 3x Gly-Ser linker.

### Secreted protein activity measurement

To measure NanoLuc activity in secretion experiments, cell cultures were separated (centrifugation 5000xg for 5-15 minutes) and supernatants were measured directly as described above or cell pellets were resuspended in 1X PBS to the volume of the initial sample before measurement. For surface anchored NanoLuc, cells first were induced with 100 ng/mL aTc and centrifuged to obtain cell pellets. Cells were then treated with 0.025% trypsin for one hour at 37C. Cells were washed twice with PBS and NanoLuc activity was measured as described.

To prepare mIL-22 samples, *C. comes* cells were grown to saturation in supplemented brain heart infusion media (BHIS: BHI + Hemin + Vitamin K_3_ + Cysteine-HCl + ATCC trace mineral supplement) containing 5 μg/mL thiamphenicol and 60 μg/mL kanamycin. Protein expression was induced by adding 100 ng/mL aTc to cells diluted 1:100 in fresh media and subsequently culturing cells at 37C for 4-6 hours to reach mid-log phase. Supernatants were collected from *C. comes* cultures *via* centrifugation for 15 minutes at 4500 rpm. Protein concentration was analyzed by mIL-22 DuoSet ELISA (R&D systems) and samples were diluted 1:100 in reagent diluent 3 (R&D systems) to avoid oversaturation of plate reader measurements.

### Bioactivity Assays

#### mIL-22 bioactivity cell culture

To measure mIL-22 bioactivity *in vitro*, HT-29MTX cells were prepared using standard culture techniques to reach 90% confluency, seeded one day prior to the assay at a concentration of 6.75×10^5^ cells per well in a 6-well plate and starved for 4-6h in DMEM (Thermo) without serum supplementation. To prepare mIL-22 samples, *C. comes* supernatants were collected from *C. comes* cultures as before and diluted in 50 μL PBS to a final concentration of 40 ng/mL secreted mIL-22 (based on concentrations measured from ELISA) or an equal volume of bacterial supernatant for the wild-type and intracellular controls. IL-22 standards (Peprotech) were prepared by dilution in PBS. Once samples and standards were prepared, they were further diluted 1:4 in DMEM without serum supplementation to a final volume of 200 μL and added to prepared HT-29MTX cells, where they were cultured for 15 minutes, washed in PBS and transferred onto ice for lysis. Cellular lysis was performed using RIPA buffer (RIPA (Thermo), micro-cOmplete protease inhibitor (Roche), HALT phosphatase inhibitor (Thermo)) on ice for 20 minutes, and cellular debris was pelleted by scraping cells into microcentrifuge tubes and centrifugation for 10 minutes at 15,000 x g. Resulting supernatant was transferred to a new microcentrifuge tube and the protein concentration was determined *via* BCA (Pierce).

Western blots of cell lysate were performed to visualize the phosphorylation of STAT3. Samples were normalized to 10 μg in Laemmli buffer containing β-mercaptoethanol and boiled for 10 minutes. Mini-PROTEAN 4-15% gels (Bio-Rad) were loaded with samples for separation at 300V for 15 minutes in Tris-SDS-glycine buffer and protein was transferred to a nitrocellulose membrane using Tris-glycine-methanol buffer for 60 minutes at 60V. Protein-containing membranes were blocked at room temperature for 2 hours in 5% milk in TBS (Thermo) with 0.05% Tween 20 (TBST). Primary incubation of 1:1000 Rb anti-Phospho-STAT3 (Tyr705 (Cell Signaling Technologies)) was conducted overnight at 4C in 5% BSA in TBST and secondary incubation of 1:5000 Goat anti-Rb IgG-HRP (7074 Cell Signaling Technologies) was conducted in 5% BSA in TBST for 1 hour. Following primary and secondary incubation, blots were washed 3 times in TBST for at least 5 minutes per wash. Blots were developed using Clarity ECL chemiluminescent substrate for 5 minutes (Bio-Rad) and imaged using a Bio-Rad Gel Doc.

Blots were stripped using Restore PLUS stripping buffer (Thermo) and washed times in TBST before re-blocking in 5% milk in TBST at room temperature for 2 hours. Blots were re-probed using Rb anti-STAT3 (D3Z2G) (Cell Signaling Technologies)) in 5% milk in TBST overnight at 4C. Secondary probing using Goat anti-Rb IgG-HRP was performed at room temperature for 1 hour and blots were washed and imaged in the same manner as for anti-Phospho-STAT3 blots.

### Mouse Experiments

#### Strains and animal procedures

All animal experiments were carried out in compliance with the University of Chicago Institutional Animal Care and Use Committee (IACUC) (Protocol 72610) guidelines and maintained at the University of Chicago Animal Resource Center (ARC). All experiments were carried out in C57BL/6J mice. Germ-free and gnotobiotic mice with a defined consortium of microbes^69^ were bred and housed within the University of Chicago Gnotobiotic Research Animal Facility (GRAF). Specific pathogen-free (SPF) mice were purchased from the Jackson Laboratory and maintained in the University of Chicago SPF facility.

Gnotobiotic C57BL/6J mice were bred and housed in Trexler-style flexible firm isolators (Class Biologically Clean) within Ancare polycarbonate mouse cages (N10HT). Gnotobiotic mice were weaned at 21 days of age and were fed an autoclaved, plant-based mouse chow (LabDiet JL Rat and Mouse/Auto 6F 5K67). All mice were housed in cages containing paper bedding (ALPHA-dri + PLUS) with a 12-hour light/dark cycle at a standard room temperature of 20-24C. All mice were euthanized by CO_2_ asphyxiation followed by cervical dislocation as a secondary measure.

### Host Intestinal Transcriptional Analysis

#### In vivo IL-22 Bioactivity in Healthy SPF Mice

SPF mice were ordered from the Jackson laboratory at 4-6 weeks of age and maintained in the facility for SPF experiments (n=4 per group). Mice were acclimated for 1 week following delivery before commencing experiments. Bacterial cells were routinely cultured (with antibiotics where appropriate) to mid-log phase as measured by microplate reader (OD_600_ of 0.4-0.6) and were diluted by 20% with 50% glycerol and either gavaged into mice immediately or frozen at −80C for future use. Each gavage consisted of 200 μL of bacteria at a concentration of 10^6^-10^9^ CFU/mL.

Bacterial dosing followed a daily, thrice weekly (MWF), or weekly gavage schedule as indicated. Following the experimental timeline, mice were sacrificed and approximately 2 cm of ileal tissue was collected roughly 1 cm proximal to the terminus of the ileum. Fatty apron was removed from the tissue section and the sample was stored in RNAlater at −80C until use.

#### RNA Preparation

RNA was extracted from ileal tissue using the RNA RNeasy PowerLyzer Tissue & Cells kit following the manufacturer protocol. RNA was normalized to 1 mg/μL in nuclease-free water. Lunascript RT SuperMix (NEB) was used for non-specific reverse transcription to cDNA and following purification (Zymo DNA Clean & Concentrator) ileal *reg3b* and *reg3g* expression was measured *via* qPCR using PowerUp SYBR Green (Thermo Fisher) following manufacturer procedures. Gene expression was normalized to *HPRT* using the ΔCt method. Alternatively, purified RNA was sequenced by Innomics using proprietary DNBSEQ technology (https://www.innomics.com/mrna-sequencing).

#### RNA-Seq Analysis

Bulk RNA-Seq reads were aligned to the *Mus musculus* reference genome (NCBI) using the STAR aligner^70^ plugin from Geneious Prime and differential expression was calculated with DeSeq2 plugin^64^ to generate count matrices for the differential expression volcano plot. Genes with adjusted p-values < 0.05 were considered significantly differentially expressed. GSEA was additionally performed using ShinyGO^71^ for STRING-db matches to GO biological processes for upregulated genes.

### Fecal 16s rRNA Analysis

At the Duchossois Family Institute at the University of Chicago Microbiome Metagenomics Facility (MMF), DNA is extracted from fecal pellets using the QIAmp PowerFecal Pro DNA kit (Qiagen) and the V4-V5 region of 16S rRNA-genes are PCR amplified using barcoded dual-index primers. Illumina compatible libraries are generated using the Qiagen QIASeq 1-step amplicon kit and sequencing is performed using Illumina MiSeq with 2×250 paired end reads (5,000-10,000 reads per sample). Raw V4-V5 16S rRNA gene sequence data is demultiplexed and processed through the DADA2 pipeline^72^ into Amplicon Sequence Variants (ASVs). ASVs are identified with the Bayesian RDP classifier^73^ up to the genus level. Linear discriminant analysis effect size (LEfSe) analysis was used to summarize taxonomic changes using default parameters^74^.

#### High Fat Diet Experiments

6-8 week old specific pathogen free (SPF) male C57BL/6 mice were purchased from Jackson Laboratory and maintained in facilities at the University of Chicago. Mice were housed in cages containing paper bedding (ALPHA-dri +PLUS) with a 12-hour light/dark cycle at a standard room temperature of 20-24°C. All mice were euthanized by CO2 asphyxiation followed by cardiac puncture as a secondary measure. All experiments were performed in accordance with the Guide for the Care and Use of Laboratory Animals (8th ed.) and were approved by the Institutional Animal Care and Use Committee of the University of Chicago (Protocol 72610).

Mice were acclimated for 1 week following delivery before commencing experiments.

Bacterial cells were routinely cultured (with antibiotics where appropriate) to mid-log phase as measured by microplate reader (OD_600_ of 0.4-0.6) and were either diluted by 20% with 50% glycerol and gavaged into mice immediately or frozen at −80C for future use. Each gavage consisted of 200 μL of bacteria at a concentration of approximately 10^8^ CFU/mL.

Western diet was produced with *ad libidum* feeding of TD.88137 (42% calories from fat) and given 0.2 μm filtered 30% fructose in the drinking water for 16-20 weeks. Mice were routinely monitored for health in accordance with IACUC guidelines. Weights were taken at the same daily time when measured.

#### Metabolic tests

Glucose tolerance tests (GTT) and fasting glucose (FG) tests were performed on 14-16 week-old SPF male mice that were fasted for 6 hours. For the GTT, mice were administered 2g/kg body weight of a 0.25g/mL D-glucose (Sigma) by intraperitoneal injection. Blood glucose was measured from a small cut to the tail vein at 0, 10, 20, 30, 60, 90, and 120 minutes using a handheld glucometer (Alphatrak3). For the insulin tolerance test (ITT), 14-16 week old mice were fasted for 2 hours. Mice were administered 1.5 U/kg of regular human insulin (Humulin-R, Ely Lilly) diluted in 0.9% saline via an intraperitoneal injection. Blood glucose was measured at time 0 (before injection of insulin) and then at 15, 30, 45, and 60 minutes after the injection of insulin.

At the end of the indicated experimental timelines, mice were weighed and sacrificed, and feces and tissue sections were collected. Livers were harvested, weighed, and sectioned for histology and downstream analyses. Blood was collected by cardiac puncture at sacrifice and centrifuged at 3,000 g for 10 minutes to isolate serum. Analyses of standard serum chemistry parameters (test code: 64014) were performed at IDEXX BioAnalytics, North Grafton, MA.

#### Liver histology

Liver sections were fixed in 10% buffered formalin, sliced into 5uM sections, and were stained with hematoxylin-eosin for pathological analysis. Scanned slides were analyzed with digital whole slide images were processed using a custom Python-based image analysis pipeline incorporating OpenCV and PIL libraries. The analysis was performed on non-overlapping 1024×1024 pixel patches extracted at 512-pixel intervals across each slide to ensure comprehensive coverage while maintaining computational efficiency. For morphological illustration, representative tissue regions were selected based on mean lipid droplet size criteria. Patches containing droplet populations with mean areas were identified and extracted as 1024×1024 pixel tiles. Each representative image was annotated with a 200 μm scale bar for standardized presentation.

Intrahepatic triglycerides (TG) were measured with a triglyceride assay kit (ab65336). Samples and standards were prepared according to the manufacturer’s instructions.

### Metabolomics

#### Metabolite Extraction from Cecal Material

Extraction solvent (80% methanol spiked with internal standards and stored at −80C) was added to pre-weighed fecal/cecal samples at a ratio of 100 mg of material/mL of extraction solvent in beadruptor tubes (Fisher). Samples were homogenized at 4C on a Bead Mill 24 Homogenizer (Fisher), set at 1.6 m/s with 6 thirty-second cycles, 5 seconds off per cycle. Samples were then centrifuged at −10C, 20,000 x g for 15 min and the supernatant was used for subsequent metabolomic analysis.

#### Metabolite Analysis

At the Duchossois Family Institute at the University of Chicago, metabolites were derivatized as described by Haak *et al*.^75^ with the following modifications.1 The metabolite extract (100 μL) was added to 100 μL of 100 mM borate buffer (pH 10)(Thermo Fisher, 28341), 400 μL of 100 mM pentafluorobenzyl bromide (Millipore Sigma; 90257) in acetonitrile (Fisher; A955-4), and 400 μL of n-hexane (Acros Organics; 160780010) in a capped mass spec autosampler vial (Microliter; 09-1200). Samples were heated in a Thermomixer C (Eppendorf) to 65C for 1 hour while shaking at 1300 rpm. After cooling to room temperature, samples were centrifuged at 4C, 2000 x g for 5 min, allowing phase separation. The hexanes phase (100 μL) (top layer) was transferred to an autosampler vial containing a glass insert and the vial was sealed. Another 100 μL of the hexanes phase was diluted with 900 μL of n-hexane in an autosampler vial.

Concentrated and dilute samples were analyzed using a GC-MS (Agilent 7890A GC system, Agilent 5975C MS detector) operating in negative chemical ionization mode, using a HP-5MSUI column (30 m x 0.25 mm, 0.25 μm; Agilent Technologies 19091S-433UI), methane as the reagent gas (99.999% pure) and 1 μL split injection (1:10 split ratio). Oven ramp parameters: 1 min hold at 60 o C, 25 o C per min up to 300C with a 2.5 min hold at 300 C. Inlet temperature was 280C and transfer line was 310 C. A 10-point calibration curve was prepared with acetate (100 mM), propionate (25 mM), butyrate (12.5 mM), and succinate (50 mM), with 9 subsequent 2x serial dilutions. Data analysis was performed using MassHunter Quantitative Analysis software (version B.10, Agilent Technologies) and confirmed by comparison to authentic standards. Normalized peak areas were calculated by dividing raw peak areas of targeted analytes by averaged raw peak areas of internal standards.

## Acknowledgments

We would like to thank the following people who help contribute to this paper: Kristen Kolar, Emma Elmiger, and Betty Theriault at University of Chicago Gnotobiotic Research Animal Facility; Alan Huff and Samuel Weng from the University of Chicago Proteomics Core; the Human Tissue Resource Center at the University of Chicago; Huaiyang Lin and Ramanujam Ramaswamy at the Duchossois Family Institute Microbiome Metagenomics Facility; Alexandra Cassano, Joyce Ghali, Jaehyun Lee, Paola Nol Bernardino, Christina Nowicki, and Sarida Pratuangtham for experimental help and insight; Matthew Brady, Eugene Chang, Nicolas Chevrier, Sam Light, Cathryn Nagler, and Eric Pamer for helpful discussions.

The authors are supported by the National Science Foundation Graduate Research Fellowship under grant no. DGE 1746045 (J.A.), by the TUBITAK BIDEB 2214-A program (R.E.A.), by the University of Chicago PREP 3R25GM066522-18 (D.V.), by the National Institutes of Health (IMSD Program 5R25GM109439-07 and Molecular and Cellular Biology training program T32 GM007183 to J.F.-S.), by the National Institute of General Medical Sciences of the National Institutes of Health (R35GM147478; M.M.), by the National Institute of Diabetes and Digestive and Kidney Diseases (P30 DK42086), and by the Arnold and Mabel Beckman Foundation through the Beckman Young Investigator Program (M.M.).

## Author Contributions

J.A., S.M., R.E.A., J.G., S.S., T.L.C., and M.M. designed experiments. J.A., S.M., R.E.A., J.G., S.S., D.V., T.L.C., J.F-S., R.M., performed experiments. J.A., S.M., R.E.A., J.G., S.S., and M.M. performed data analysis. J.A., S.M., and M.M. conceived this study and wrote the manuscript.

## Competing Interests

A patent (PCT/US2024/031745) related to this research has been filed by The University of Chicago, with M.M., J.A., and R.E.A. as inventors.

