## Supplemental Data for "Engineering butyrate-producing Lachnospiraceae to treat metabolic disease"

### Supplemental Figure 1

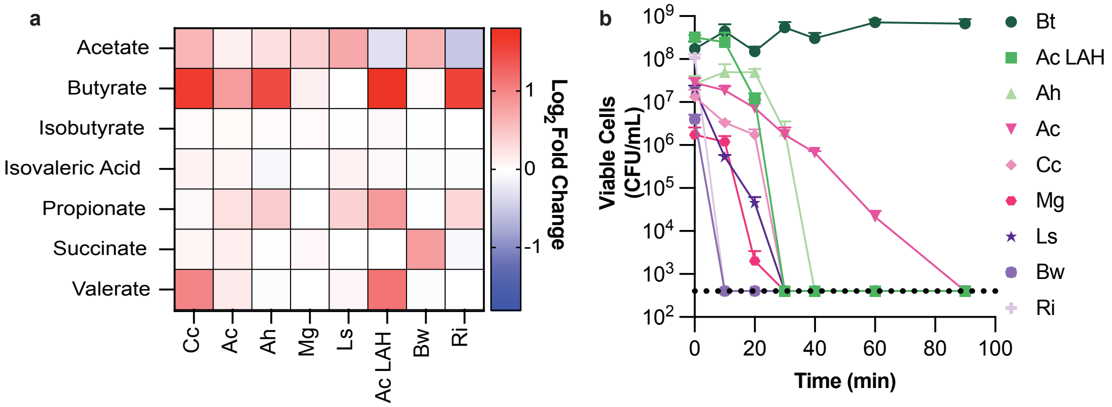


**Supplementary Figure 1.**

(a) Short-chain fatty acid quantification of Lachnospiraceae species *via* PFBBr derivatization (*Coprococcus comes ATCC 27758,* Cc; *Anaerostipes caccae DSM 14662,* Ac; *Anaerostipes hadrus DSM 3319,* Ah; *Mediterranibacter gnavus ATCC 29149,* Mg; *Lachnoslostridum symbiosum WAL 14163,* Ls; *Anaerostipes caccae LAHUC,* Ac LAH; *Blautia wexlerae DSM 19850*, Bw; *Roseburia intestinalis DSM 16841*, Ri).

(b) Relative aerotolerance test of Lachnospiraceae species (*Bacteroides thetaiotaomicron ATCC 29148*, Bt – aerotolerant control).

### Supplemental Figure 2

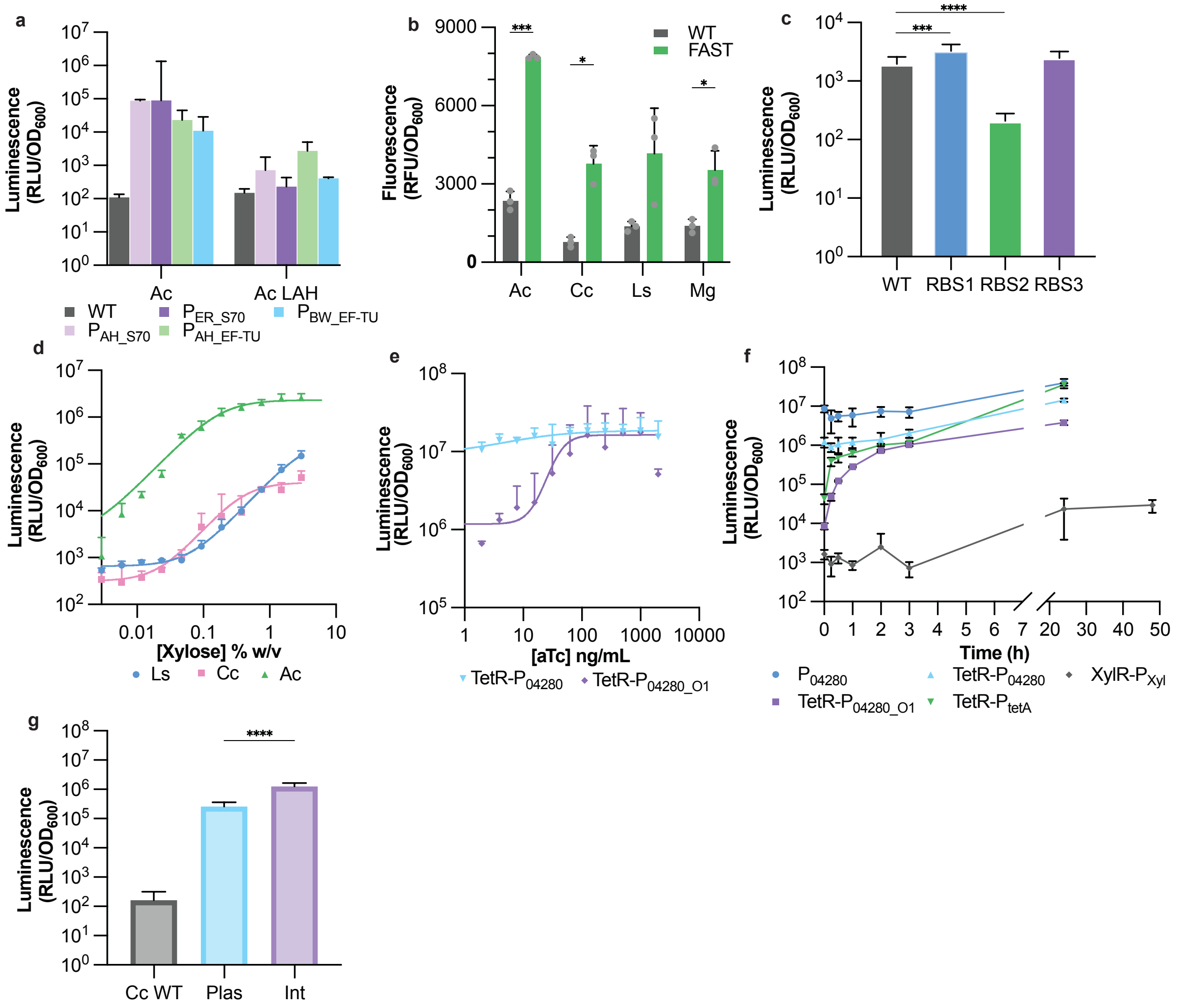


**Supplementary Figure 2.**

(a) Promoter screen in *Ac* using the wild-type strain (Ac WT) and a laboratory isolate from UChicago (Ac LAH).

(b) Comparison of Fluorescence-Activating and Absorption-Shifting Tag (FAST) versus wild-type (WT) activity in four species (Ac, Cc, Ls, Mg). All conditions were treated with 20 µM HMBR. *, P < 0.05; ***, P < 0.001 (T-test).

(c) Evaluation of three synthetic ribosome-binding sites (RBS1–RBS3) driving NL activity in Cc with a minimal promoter based on $P_{4280}$. The minimal promoter motif was predicted by promoter-id-from-rna-seq[48]. The predicted -35 site is TTGACT and the predicted -10 site is TATAAT. ***, P < 0.0005; ****, P < 0.0001 determined by one-way ANOVA.

(d) Dose–response of xylose-regulated NL expression in Ls, Cc, and Ac.

(e) Schematic of synthetic aTc-inducible promoter and aTc-dependent dose-response NL in *Cc comparing the synthetic inducible promoter (*TetR-$P_{04280\_O1}$) and an operator free promoter (TetR-$P_{04280}$).

(f) Kinetic profiling was performed for different inducible promoters in *C. comes*. Inducer was added to mid-log phase cultures at a concentration of 1000 ng/mL for [aTc] or 1.5 % w/v for [Xyl]. $P_{04280}$ was used as a positive control.

(g) Initial NanoLuc activity for plasmid maintenance assay (Fig. 1f-g). *, $P < 0.05$; ****, $P < 0.0001$ determined by one-way ANOVA.

Bars/points show geometric mean ± geometric s.d.

### Supplemental Figure 3

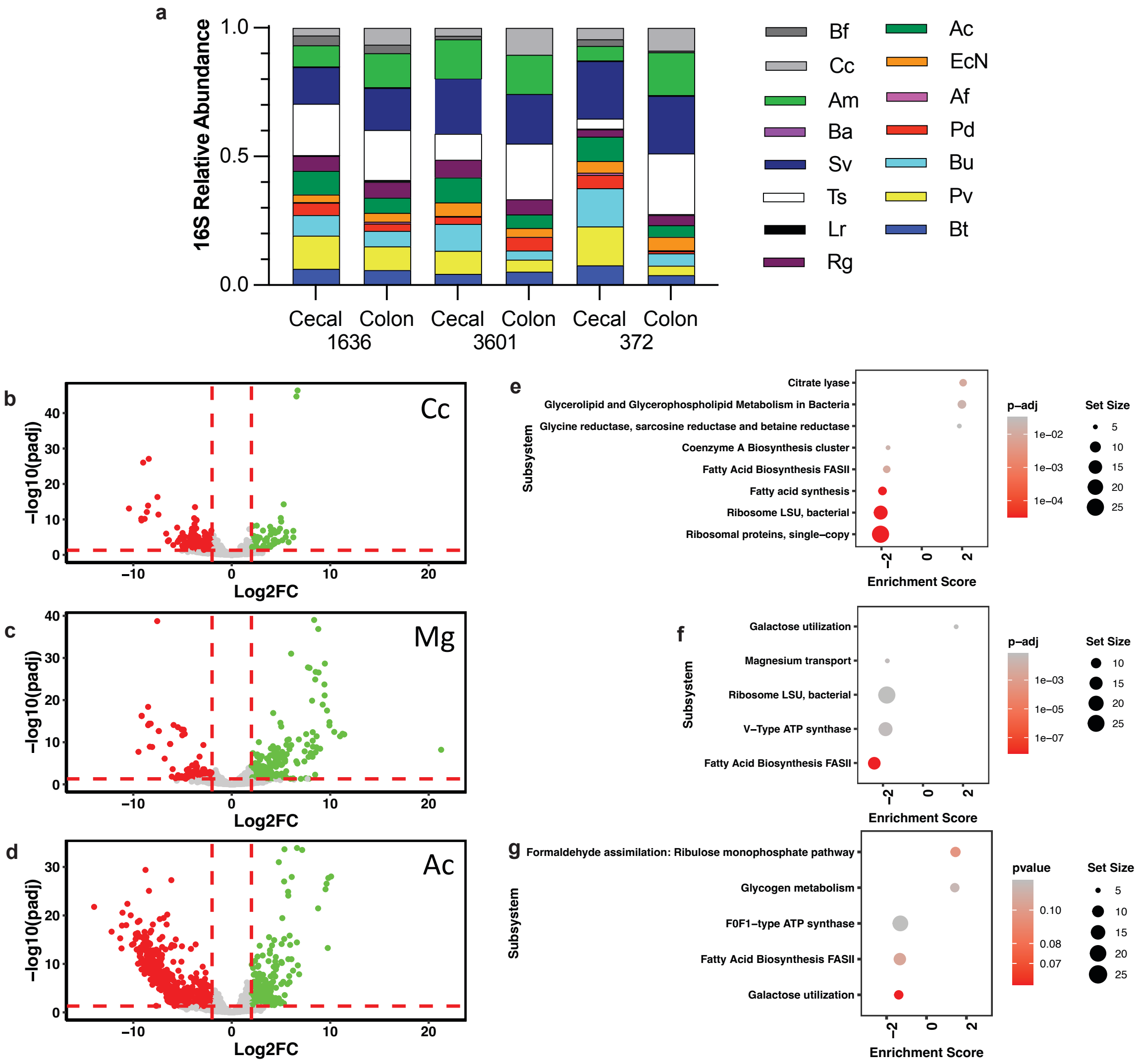


**Supplementary Figure 3.**

(g) Relative abundances of bacterial taxa in fecal and cecal samples from defined-community-colonized mice, providing the compositional background for in vivo transcriptomic profiling.

(b, c, d) Volcano plot of in vitro vs in vivo differential expression analysis of Coprococcus comes (Cc), Mediterraneibacter gnavus (Mg), and Anaerostipes Caccae (Ac). Genes that are significantly up- or downregulated genes (Log2FC > 2 or Log2FC < -2, p-adj < 0.05) are marked by color (green and red).

(e, f, g) Corresponding dot plots represent Gene Set Enrichment Analysis (GSEA) performed on expression analysis using clusterProfiler. Data shown are SEED Subsystems that were significantly enriched (p < 0.05) for each species.

**Supplemental Figure 4**

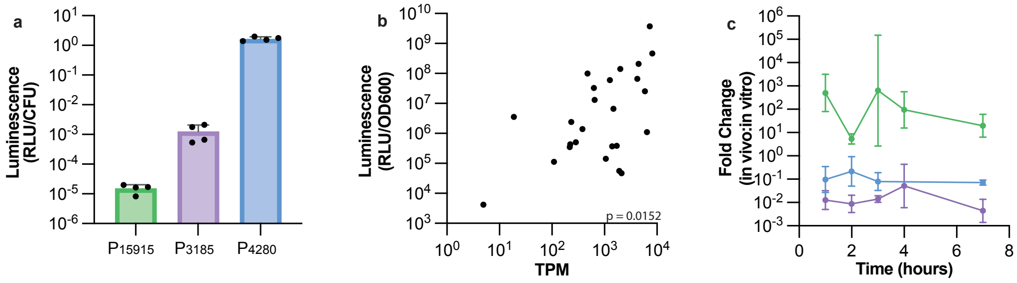


**Supplementary Figure 4.**

(a) *In vitro* expression of selected promoters from the RNA-seq promoter library used for *in vivo* validation. $P_{15915}$ (enriched *in vivo*), $P_{3185}$ (enriched *in vitro*), and $P_{4280}$ (highly expressed in both environmental conditions).

(b) Correlation plot of Promoter Luminescence (RLU/OD600) at mid-log growth vs Transcripts per Million (TPM) measurement from RNA-seq. Spearman's rho was calculated in Prism, demonstrating a modest correlation between the two values.

(c) Fold change from *in vivo* and *in vitro* experiments to compare promoter expression (RLU) per Colony Forming Unit (CFU) at measured time points. Fold Change calculated as *in vivo* over *in vitro*.

### Supplemental Figure 5

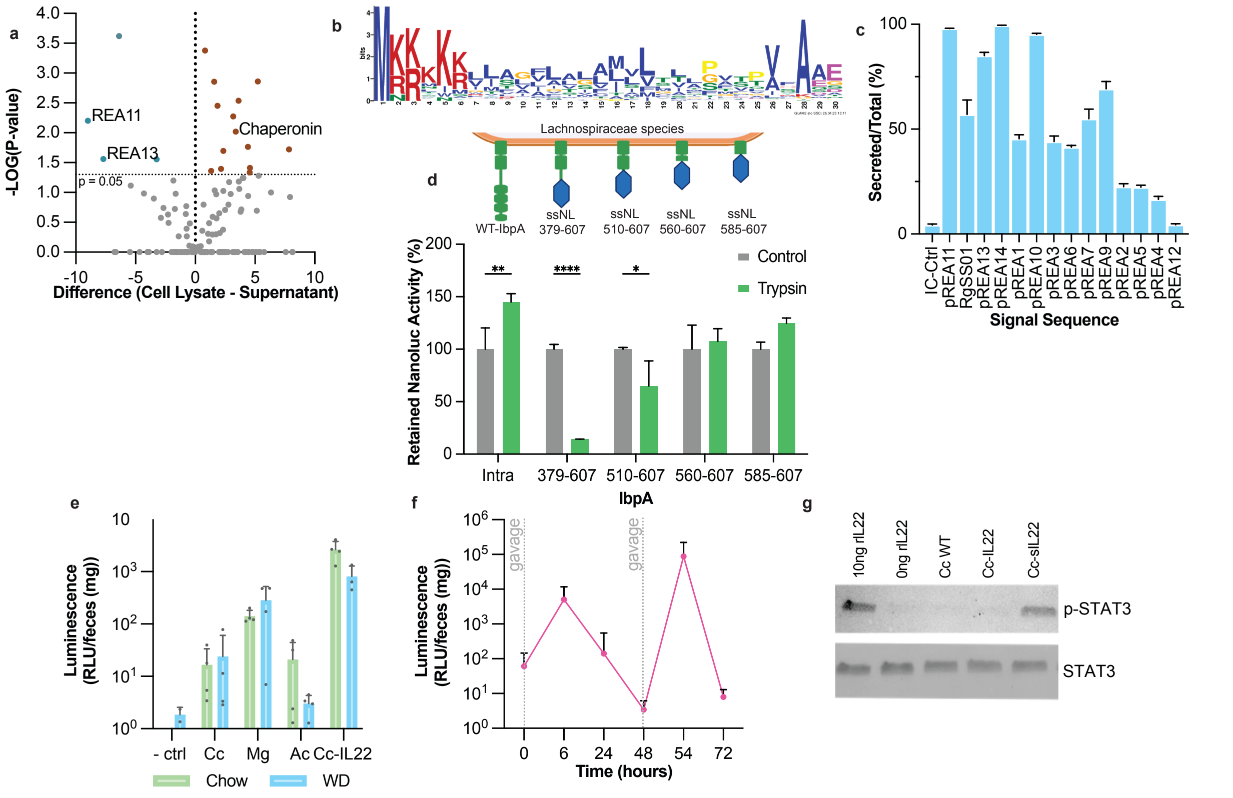


**Supplementary Figure 5.**

(a) Volcano plot of identified proteins from cellular and extracellular fractions of *in vitro* Mg cultures. REA11 and REA13 were identified from this screen as indicated on the plot (blue circles). GroEL chaperone is identified as a proxy for cytosolic proteins.

(b) Synthetic signal sequence motif, REA14 generated using GLAM2[71,72].

(c) Secretion efficiency was determined as the ratio of NanoLuc activity detected in the supernatant fraction compared to total NanoLuc activity in the *in vitro* Cc culture (cell pellet and supernatant).

(d) Schematic of surface anchoring motif discovery from *IbpA* truncations. NanoLuc activity measured following trypsin treatment.

(e) Fecal luminescence measured 6 h after oral gavage of NanoLuc-expressing strains Cc, Mg, Ac, and Cc-IL22 in chow or HFD fed mice. (n=4 per group)

(f) Fecal luminescence following repeated gavage of NanoLuc-expressing Cc-IL22. (n=10)

(g) Immunoblot for phosphorylated STAT3 in HT29-MTx cells stimulated with recombinant murine IL-22 (mIL-22) or with supernatant from Cc-IL22 or Cc culture respectively. Recombinant murine IL-22 (Peprotech) dissolved in PBS was used as a positive control.

Bars represent mean ± s.d. Statistical significance determined by two-way ANOVA: *, $P < 0.05$; **, $P < 0.005$; ****, $P < 0.0001$.

### Supplemental Figure 6

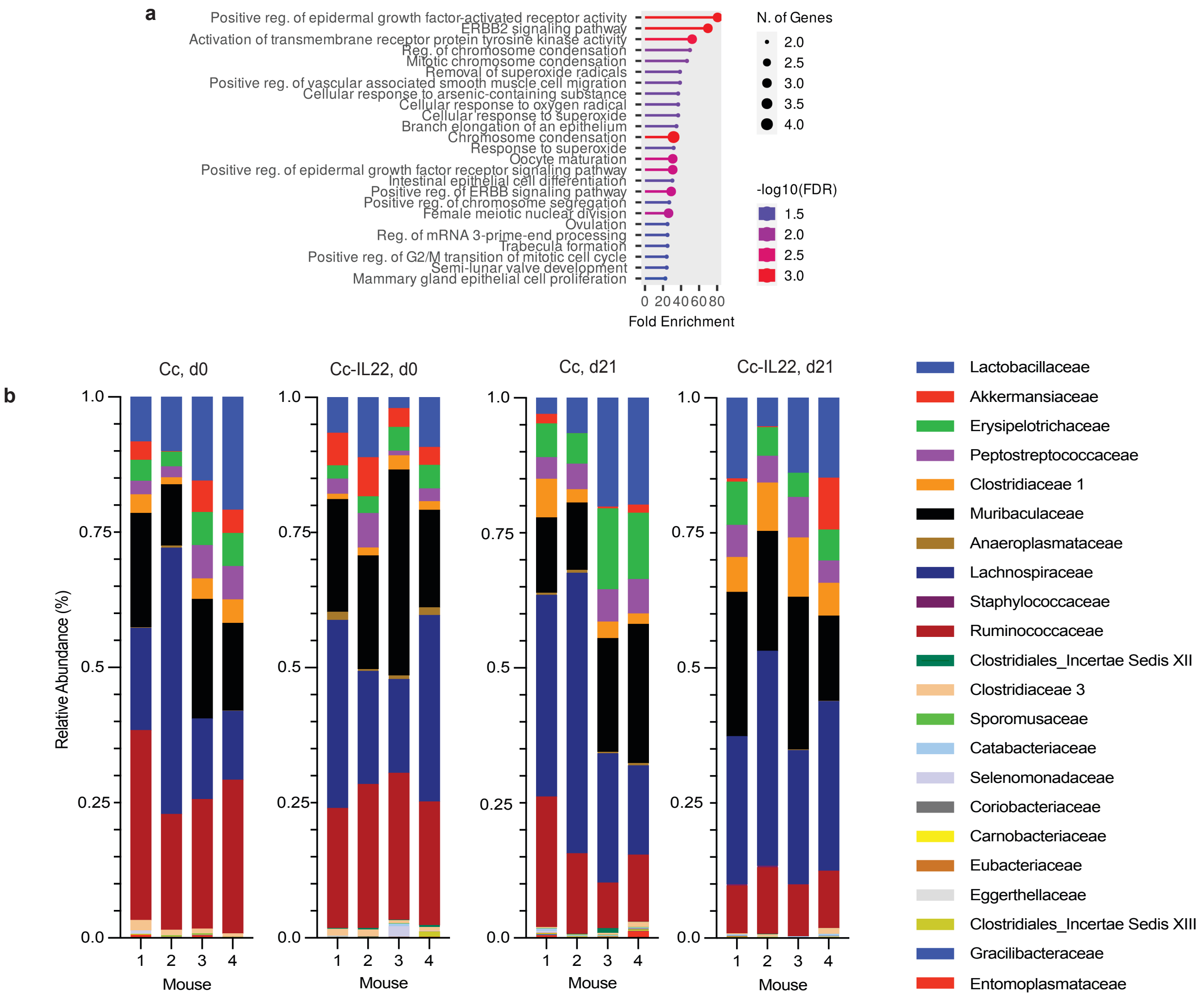


### Supplementary Figure 6.

(a) Gene set enrichment of annotated GO terms from murine ileal tissue for chow-fed mice treated with Cc-IL22 compared to mice treated with Cc after 21 days using ShinyGO[73].

(b) Relative abundances in fecal samples measured by 16S rRNA gene sequencing at baseline and after 21 days in chow-fed mice given Cc or Cc-IL22 (*n* = 4 per group). Bars show per-sample family-level composition.

### Supplemental Figure 7

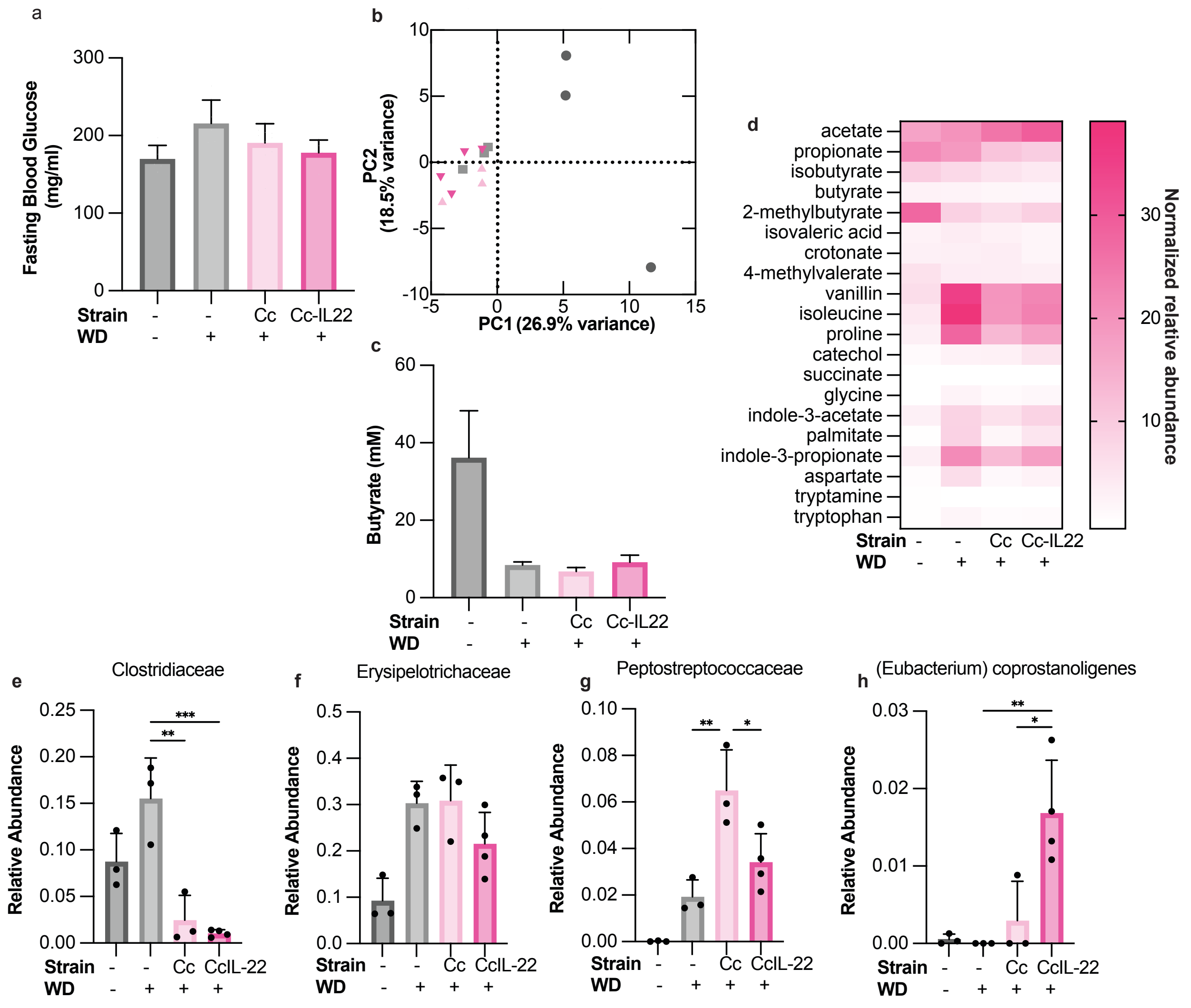


**Supplementary Figure 7.**

(a) Fasting blood glucose levels at 14 weeks.

(b) PCA plot of 16S rRNA gene sequencing data at the family level.

(c) Cecal butyrate levels at 16 weeks.

(d) Cecal short-chain fatty acid (SCFA) profiles determined by metabolomics.

(e, f, g, h) Relative abundance of bacterial families quantified by 16S rRNA sequencing, with families showing significant differential representation indicated.

Error bars represent mean ± s.d. Statistical significance determined by one-way ANOVA:*, P < 0.05; **, P < 0.01; ***, P<0.001.

### Supplemental Figure 8

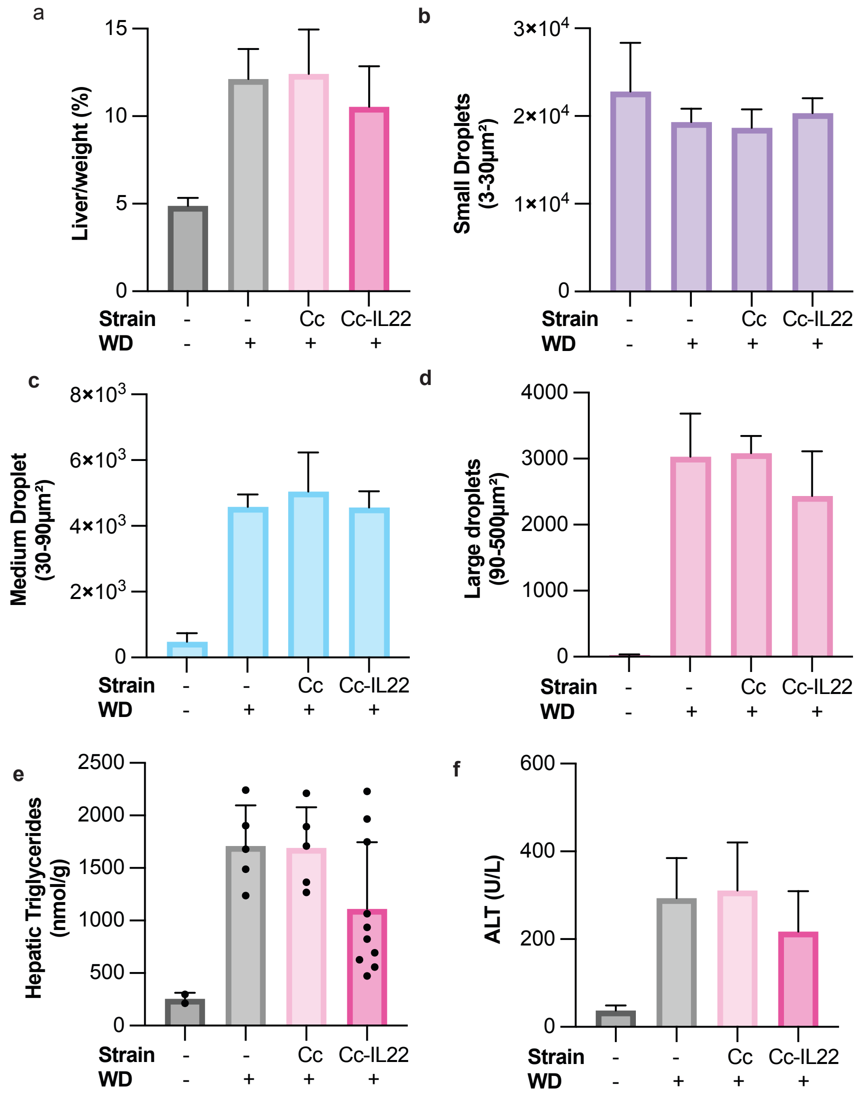


**Supplementary Figure 8.**

(a) Percent liver weight at 16 weeks, calculated as liver weight relative to total body weight.

(b, c, d) Quantification of small, medium, and large droplets present across 100 full tissue patches per sample.

(e) Hepatic triglyceride concentrations.

(f) Serum alanine aminotransferase (ALT) levels.

Error bars represent mean ± s.d.
